# Control of density and composition in an engineered two-member bacterial community

**DOI:** 10.1101/632174

**Authors:** Reed D. McCardell, Ayush Pandey, Richard M. Murray

## Abstract

As studies continue to demonstrate how our health is related to the status of our various commensal microbiomes, synthetic biologists are developing tools and approaches to control these microbiomes and stabilize healthy states or remediate unhealthy ones. Building on previous work to control bacterial communities, we have constructed a synthetic two-member bacterial consortium engineered to reach population density and composition steady states set by inducer inputs. We detail a screening strategy to search functional parameter space in this high-complexity genetic circuit as well as initial testing of a functional two-member circuit.

We demonstrate non-independent changes in total population density and composition steady states with a limited set of varying inducer concentrations. After a dilution to perturb the system from its steady state, density and composition steady states are not regained. Modeling and simulation suggest a need for increased degradation of intercellular signals to improve circuit performance. Future experiments will implement increased signal degradation and investigate the robustness of control of each characteristic to perturbations from steady states.

## I. INTRODUCTION

Microbiome research is diving deeper and deeper into microbial communities and continues to uncover new important roles for humans’ resident microbes in normal physiology. Recent work has provided mechanistic explanations for previously reported correlations between microbiome composition and disease states [1–4]; limited attempts to therapeutically alter or supplement microbiome composition have proven highly effective for altering microbiomes and treating disease [5, 6]. Clearly, it is in our interest to understand mechanisms that can control the composition of our microbiomes so we might precisely engineer them for health in the future.

Bioengineers have recognized the need for improved measurement and manipulation of hard-to-access microbiomes and produced an array of technology that gives us a window into our resident microbial world and improved ways of interacting with it [7–9]. With the ability to measure and affect our microbial communities, it is important to determine the right way to affect a microbiome to produce a desired change. Biologists and control engineers have tackled this question and produced significant theoretical guideposts for experimental work with microbial communities [10, 11] as well as important fundamental experimental results in simple microbial systems.

One of the earliest experimental demonstrations of microbial community control combined two fundamental technologies mined from microbial physiology [12]. The ccdB toxin and acyl-homoserinelactone (AHL) quorum sensing were used to construct a simple feedback system for control of a monoculture’s population size. Later work scaled up to multi-membered systems using similar pieces of genetic technology. Balagaddé *et al* used ccdA, the ccdB antotixin, to design a circuit recapitulating the out of phase oscillations of a predator-prey ecology [13]. Scott *et al* replaced ccdB with the Φ*X*174 phage lysis gene in a system of two non-interacting self-limiting bacterial populations that maintain a heterogeneous community, rather than collapsing to monoculture, due to oscillations in each member’s density [14]. These systems are important explorations into synthetic circuit design for the regulation of two-membered communities. We hope to use this library of genetic technology to address stable feedback control of multiple characteristics of heterogenous community, expanding on the original demonstration of feedback control in a monoculture.

We have previously reported a genetic circuit motif for the construction of multifunctional bacterial community controllers [15]. Using the ccdB/A toxin antitoxin system and AHL quorum sensing, the circuit motif can implement pseudo-integral control [16] and can be easily engineered to take arbitrary inputs for different control circuit architectures.

## II. Results

Symmetrically connecting our previously reported circuit motif in two different cells produces a community circuit that can control multiple characteristics of the population (Fig. 1). Two inducers activate AHL production in each cell, signaling negative feedback for each producer and rescue for the partner. Each cell is labeled with a constitutively expressed fluorescent protein to allow quantification of the abundance of each cell type in the community. Initial simulations of circuit function showed tight control of population density, but poor control of population composition, which appeared to correct past its set point and stabilize at a point dependent on initial community conditions once the population steady state was achieved. This dependence on initial conditions was also observed when solving for density and composition steady states analytically. Nevertheless, we built the circuit while debugging its design to see if it would perform in experiments as it did in simulations.

**Fig. 1.**
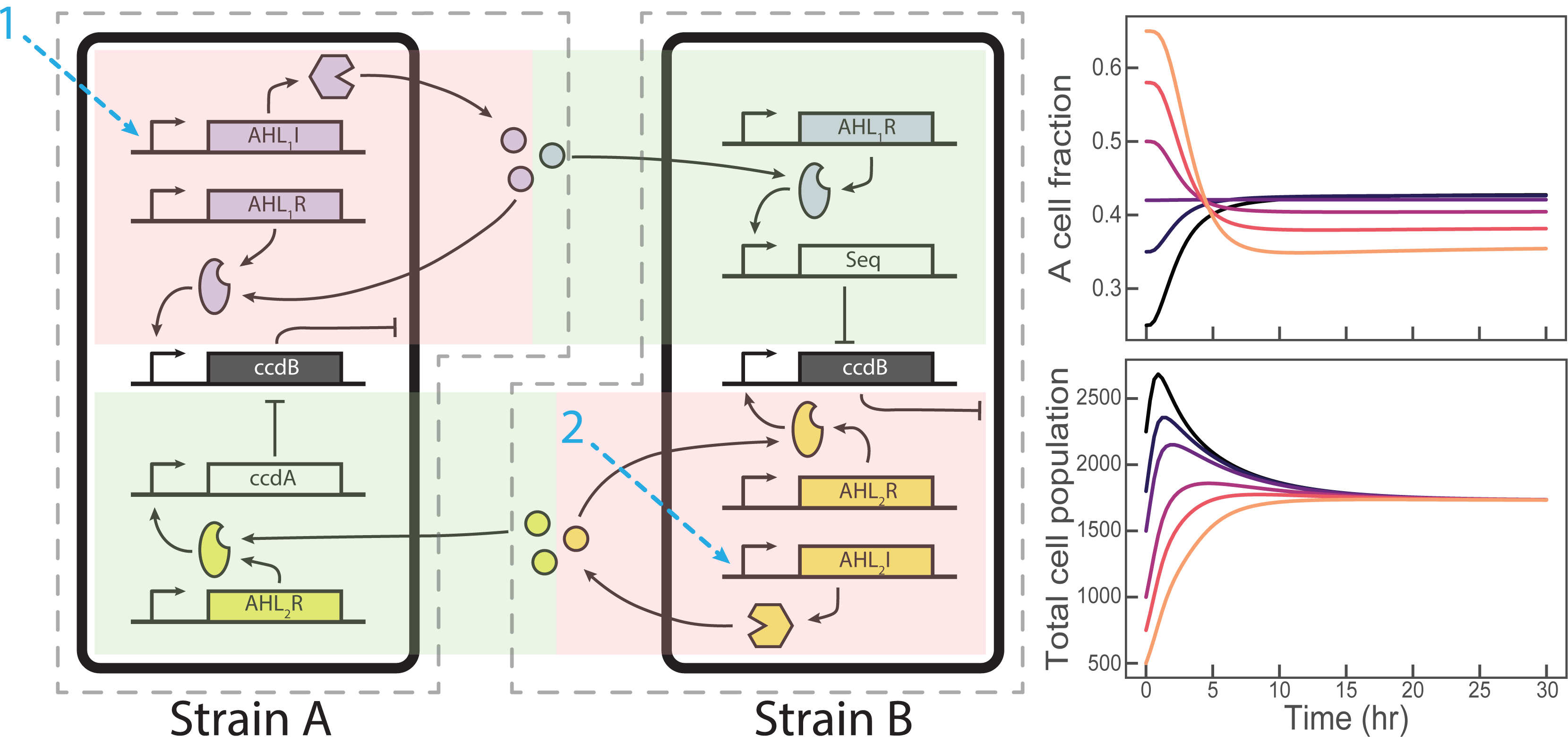
The A=B circuit uses a symmetric circuit motif in its two cells to create cis-acting negative feedback loops on each member and trans acting rescues from negative feedback from each member to the other. Negative feedback and sequestration rescue are effected by ccdB toxin and its ccdA antitoxin, respectively. Signals 1 and 2 are chemical inducers that activate transcription of AHL synthase genes. Plots are simulations of cell density and composition. Each colored curve represents seeding a community at a unique composition and density. Without intercellular signal degradation, density converges to a steady state while composition does not converge.

The ccdB protein is a highly potent toxin and slight overexpression can very easily lead to total population death. As such, the parameter ranges in which this circuit design is actually a functional controller are tight. Sensitivity analysis [17, 18] of a mathematical model of the circuit shows that circuit function is sensitive to a few of the parameters in the model, especially the ccdB expression rate (Fig. 6). Thus, to create the functional circuit in the laboratory we chose to screen circuit variants with different ribosome binding site (RBS) strengths driving ccdB translation to search the widest range of circuit functional space. 3G assembly [19] was used to assemble the genetic parts of this circuit using a pool of different-strength RBS’s driving translation of the ccdB toxin in each cell’s motif. All other circuit components were designed to be expressed with hardcoded intermediate strength.

Assembly and screening of the different ccdB expressing variants of each cell had 2 selection phases, a negative selection against cells overexpressing ccdB (these did not grow colonies for selection during cloning) and a functional screen for appropriate community behavior in a simple experiment observing coculture behavior with the circuit “ON” (both inducer inputs at maximal induction) and “OFF” (no inducers present). Total density was measured with absorbance at 700nm and composition was measured with flow cytometry.

Communities displaying reductions in steady state total population density and changes in steady state population composition (without collapsing immediately to monoculture) were considered candidate functional circuit designs.

Consistent with model predictions, all screened communities of A and B cell variants displayed population control behavior with maximal inducer concentrations (Fig. 2). Composition control was more sensitive to the variations in each culture. Cell variant mixtures were sampled at different times during the screen and measured with flow cytometry to quantify the number of cells of each type. Some cocultures did not appear to achieve different compositions in different inducer conditions, others quickly collapsed to monocultures of just A or B cells. A minority of cocultures maintained a mixed composition that was modified by inducer concentrations. These cocultures were considered candidate functional cell mixtures and saved for more detailed experimental testing.

**Fig. 2.**
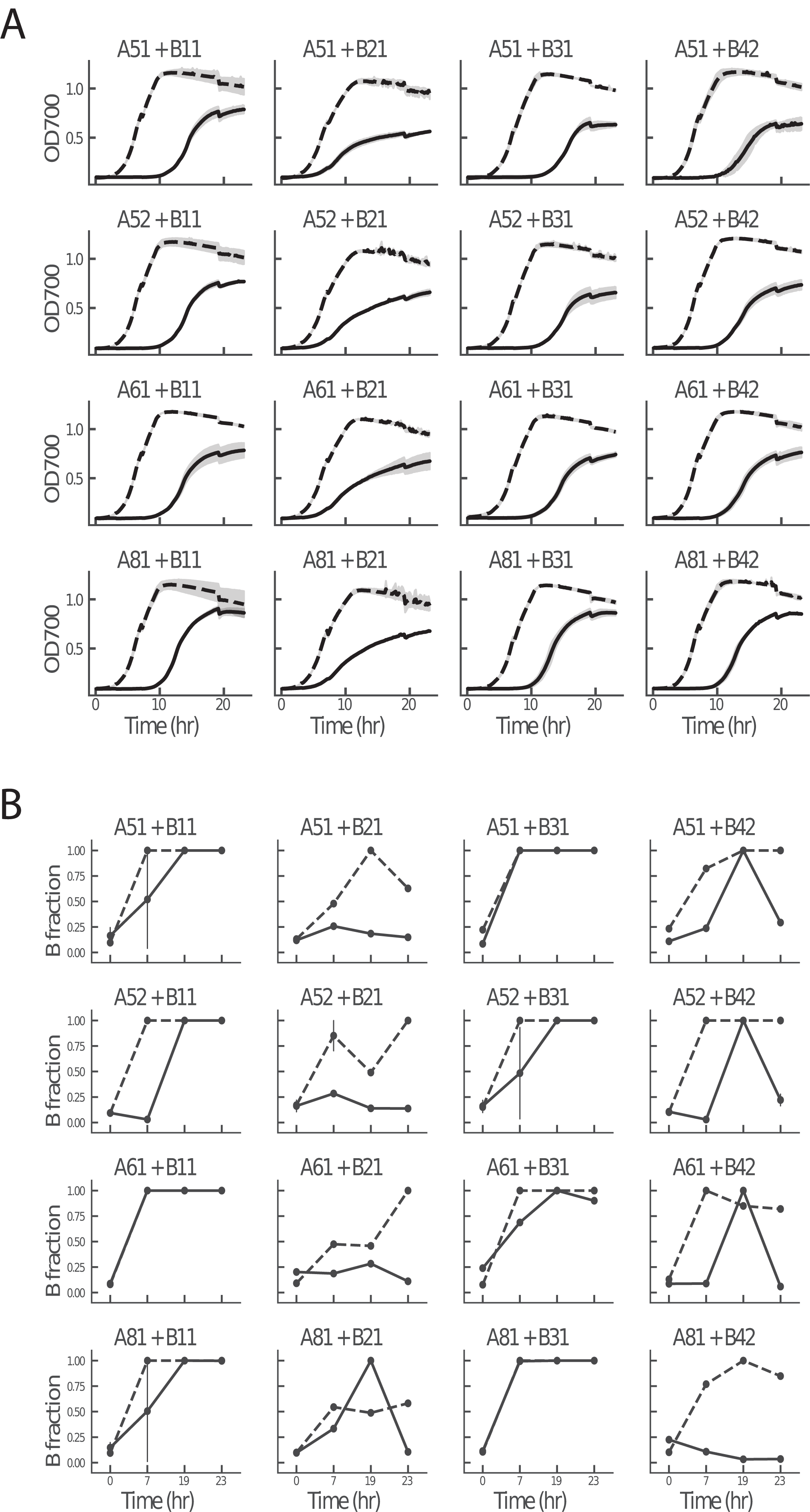
Mixtures of variants of each cell type scan parameter space, demonstrating that total density control is much more robust to parameter changes than composition control. Induced mixtures of cells are drawn as solid lines, uninduced mixtures are dashed. (A) Total community density measured by OD700 is capped at a varying steady states at maximal circuit induction, depending on the mixture of A and B cell variants. (B) Calculating the fraction of B cells in the community from flow cytometry reveals some communities collapse quickly to monocultures and/or show no significant response to inducer changes, which others maintain mixed cultures whose compositions change with inducer changes.

The first design iteration through this screening procedure used a DH5*α*-Z1 derived *E. coli* strain integrated in-house with the CinR and RpaR AHL activator proteins. This strain did not reliably maintain fluorescent label expression in experiments longer than 24 hours, so the base strain was switched to the Marionette-wild type *E. coli* developed by Meyer *et al* [20] and the AHL synthases/promoters were replaced with the Lux and Cin sets to make use of the Marionette strain’s well characterized receptor modules. The same assembly and screening method using the Marionette cells and the updated circuit plasmids generated a few candidate A and B cell variants that were used in later experiments.

We hoped to demonstrate tunable population density and composition steady states by changing inducer concentrations. Candidate cell mixtures were grown in 4 inducer conditions in which both inducer concentrations were increased together. The cultures were grown for 18 hours, then diluted 1:10 in identical inducer concentrations to test the system’s robustness to perturbation. Different inducer concentrations set multiple density and composition steady states, but circuit function appeared limited to one growth phase (Fig. 3).

**Fig. 3.**
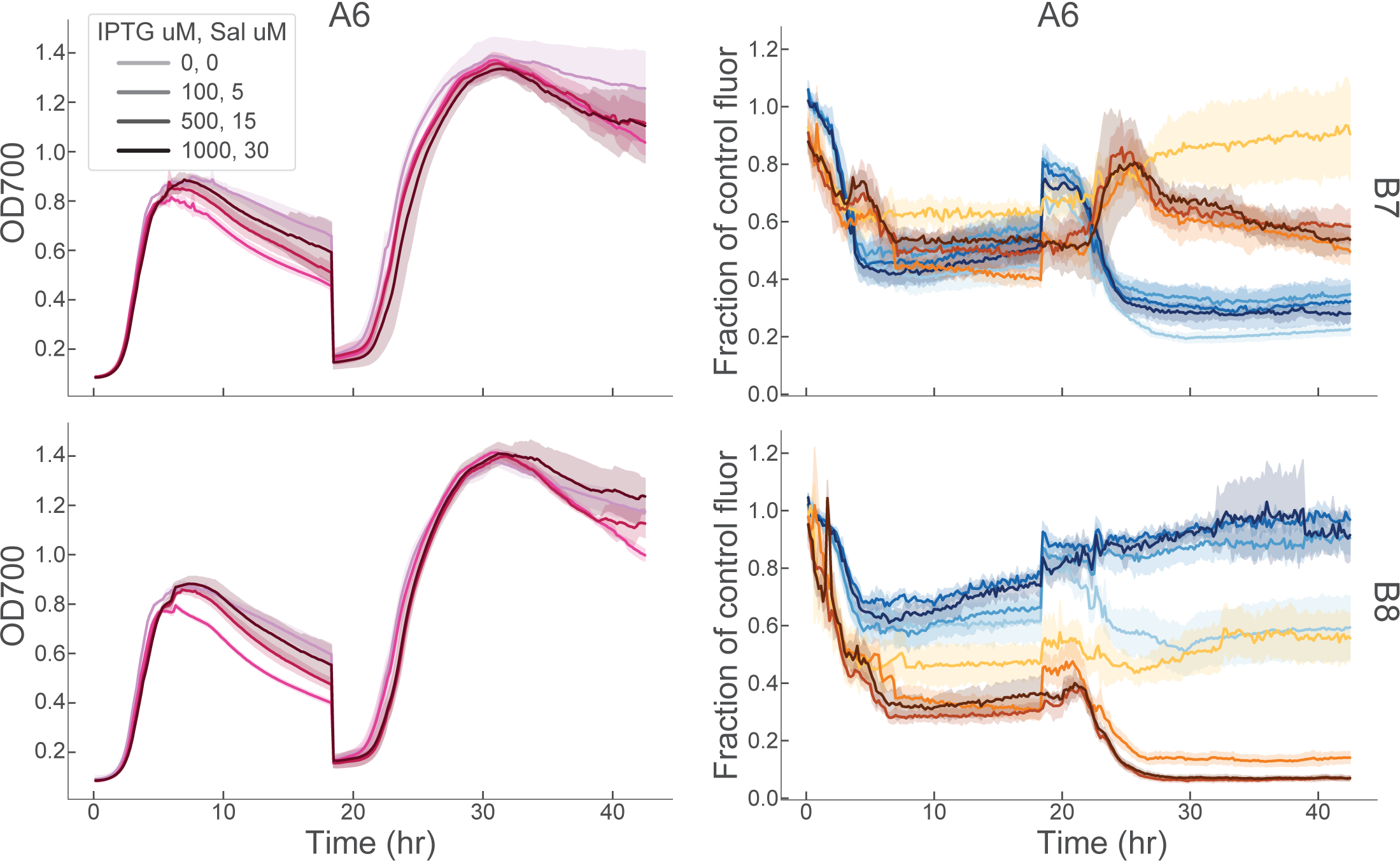
Two community variants that passed density/composition control screening are composed of A and B cell variants A6, B7, B8 in all AB combinations. (*Right*) These mixtures respond to inducer by setting different density and composition steady states. Density control is much less noticeable in these variants; the base *E. coli* strain was altered to improve fluorescence intensity. (*Left*) Comparing fluorescence intensity between experimental mixtures and monocultures of each A or B cell in identical inducer concentrations, we can estimate population composition. Cultures were diluted 10x at 18 hours and grown to steady state a second time.

Population density was capped during the first growth phase in all cases at or below OD 1.0. In the second growth phase, density control appears to be lost, the cultures grew to the carrying capacity of the vessel (OD700 1.4). This could be a consequence of circuit breakage due to mutation or other escape in one or both of the cells. OD700 measurements alone were insufficient to diagnose where the circuit problem occurred, fluorescence measurements that track the abundance of each cell type provide more insight.

Fluorescence measurements of experimental mixtures and A or B cell monocultures were compared to estimate population composition. Uninduced mixtures maintained an approximately 1:1 population composition; one mixture was unperturbed by dilution, while the other was destabilized and collapsed to a B cell monoculture. Increasing inducer concentrations did not affect the A6+B7 mixture in the first growth phase, but prevented collapse to B cell monoculture in the second growth phase, consistent with at least partial function of the circuit. Induction of A6+B8 affected a mild bias towards predominance of the CFP labeled A cells in the first growth phase, which was only amplified in the second. The concurrent loss of density control in both mixtures can explain the frequent collapses to monocultures, suggesting that one or both of the cells in the mixture escape regulation by their circuit motifs and grow to the carrying capacity of the vessel (Fig. 3).

It is important to remember that bulk fluorescence and OD700 are not direct measurements of the abundance of living cells of each type. In the past, we found OD700 to significantly overestimate the number of viable cells in culture when the culture was expressing ccdB toxin to cap its density. ccdB does not lyse cells when it kills them, leaving dead cells that still absorb at 700nm and may still contain actively fluorescent proteins, complicating the interpretation of our absorbance and fluorescence measurements. To directly measure both the population density and composition, we counted viable cell numbers at different times in the experiment, using a colony forming units (CFU) assay.

Where OD700 did not detect a difference in total population density in the second growth phase in any of the inducer conditions, CFU counts revealed significant continuing effects of inducer on total population density, though the steady states from the first growth phase were not maintained (Fig. 4). In addition, increasing total induction produced expected, large decreases in population density, rather than the small changes observed with OD700 absorbance. CFU counts of each cell type agreed with the trends observed in bulk fluorescence, showing collapse of the A6+B7 population to an B cell monoculture and the opposite in the A6+B8 culture. CFU counting is the most accurate method for measuring population density and composition in our system, but bulk fluorescence is also valuable for its high time resolution and good concordance with CFU data.

**Fig. 4.**
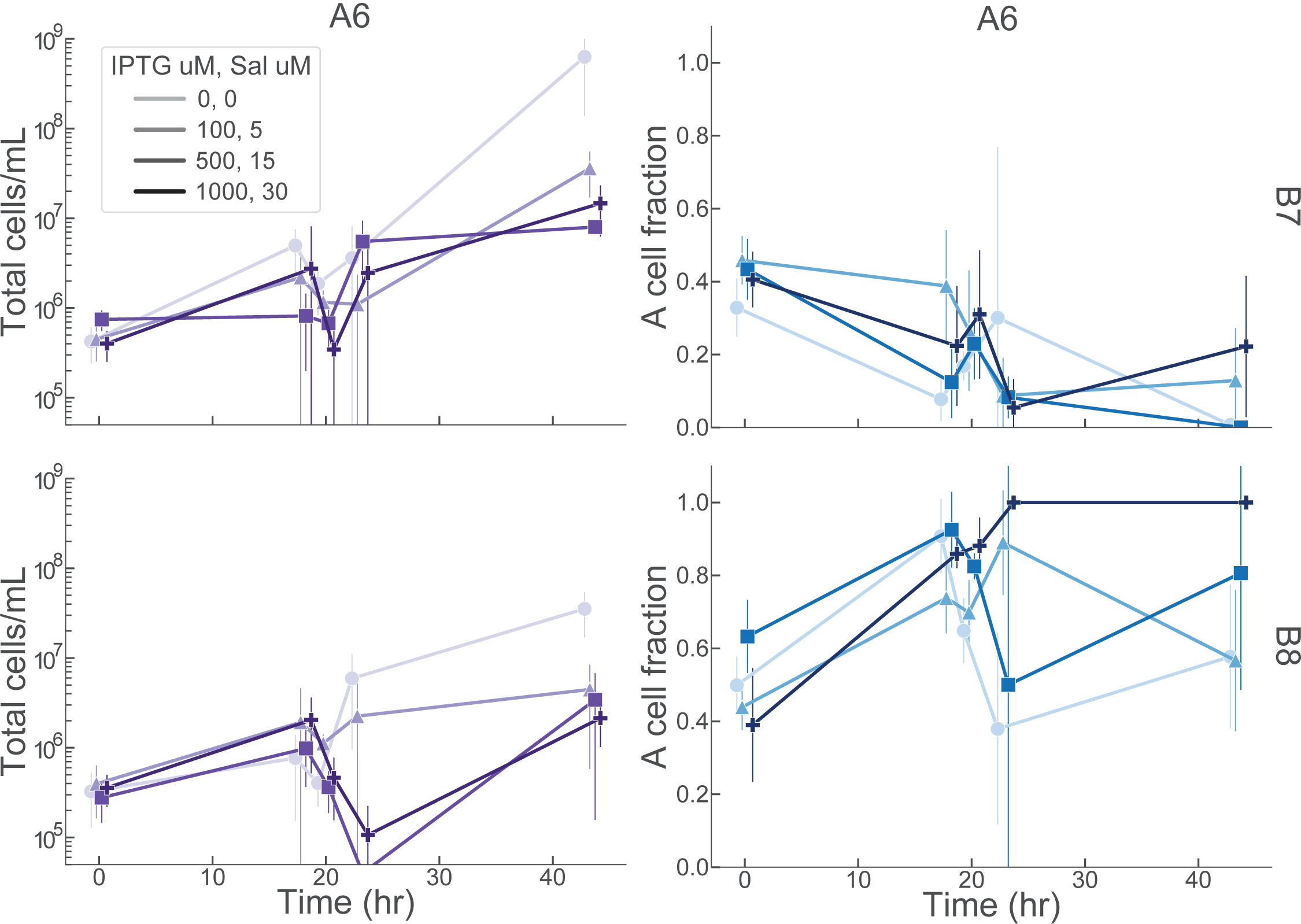
The mixtures of variants A6, B7, B8 were plated for CFU counting at t = 0, 18, 18 (post dilution), 23, 43.5 hours. Each set of points is offset slightly on the time axis to avoid obscuring data. (*Left*) CFU counting reveals inducer effects on population density hidden by OD700 measurements. (*Right*) Counts of each cell type agree with composition trends observed with bulk fluorescence. Error bars are very wide due to false negative colony counts in which a culture grew zero colonies at a particular dilution. This is an experimental artifact that will be resolved with more sampling at a wider range of dilution factors.

## III. DISCUSSION

We set out to build a population level genetic circuit for the control of population density and composition using the toolbox of biological parts useful for engineering bacterial communities. Initial tests of community variants showed obvious effects of circuit induction on both population density and composition, but both simulation and experimentation revealed a lack of steady state stability and significant sensitivity to circuit parameter changes. In this state, our design could not be called a true controller of community characteristics.

After revisiting our design, we realized our system allowed the production of AHL signaling molecules without bound, far into saturating AHL concentrations that prevent establishment of composition steady states (imperfect composition control in simulation) and perturbation rejection (inability to correct after a dilution). Most studies of AHL-linked bacterial populations employ active AHL degradation or dilution via enzymatic degradation or dilution in a chemostat. We had overlooked this critical component of our design and observed the appropriate, if undesired, lack of circuit function in simulations and experiments. Simulations including active AHL signal degradation drastically improve the accuracy of the controller and robustness to perturbation (Fig. 5).

**Fig. 5.**
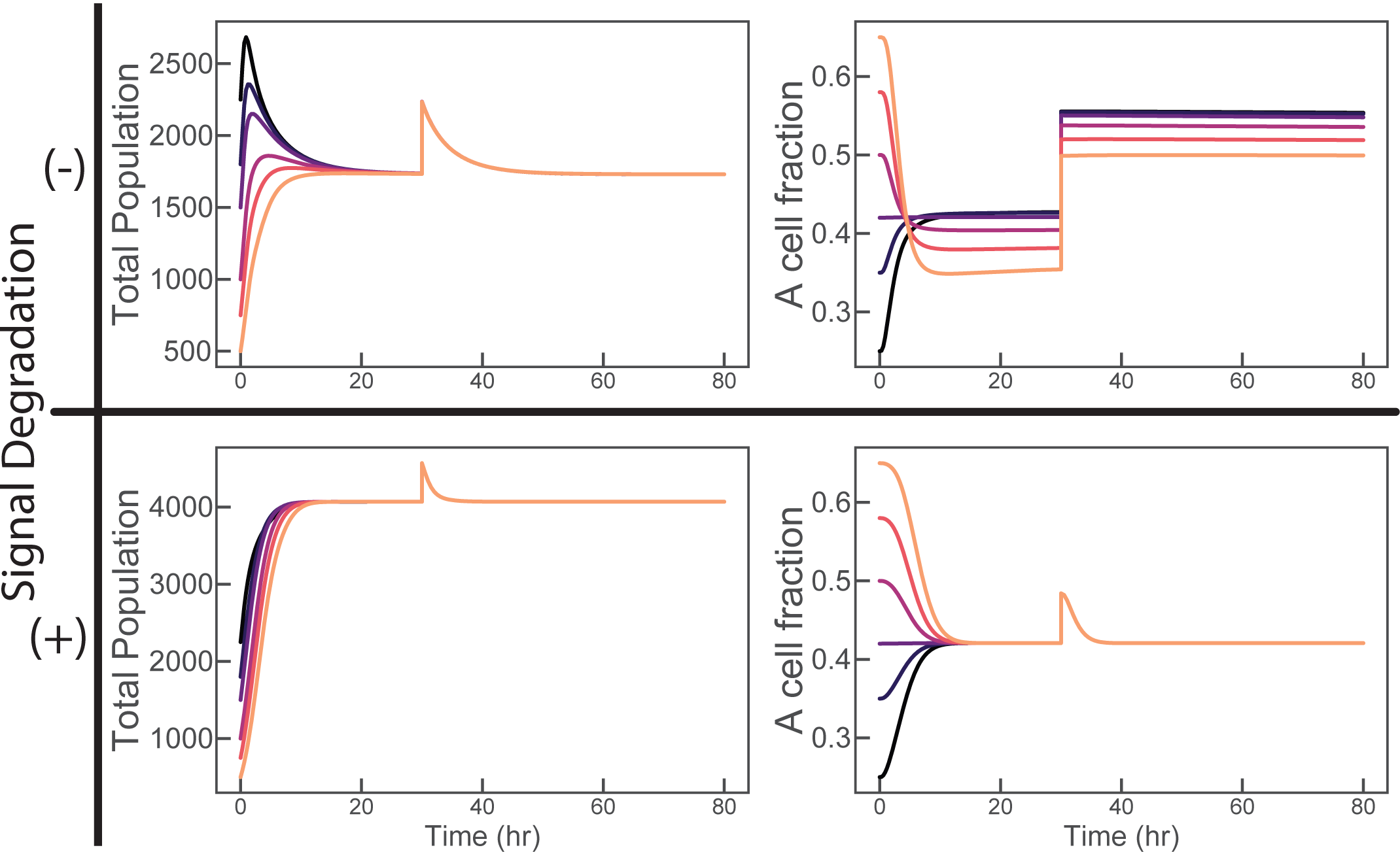
Plots are simulations of cell density and composition. Each colored curve represents seeding a community at a unique composition and density. At t = 30hr, 500 A cells are added to the community to perturb the steady state. Without intercellular signal degradation, density converges to a steady state and can reject the perturbation; composition does not initially converge and does not respond at all to perturbation. With degradation, density converges to a higher steady state consistent with lower AHL signal concentrations and rejects the perturbation; composition both converges initially and can reject the perturbation.

We encountered a number of experimental trade-offs throughout testing our circuit. In previous work, we learned that ccdB’s toxic action is not accurately captured with cell absorbance measurements. Cells killed by ccdB are not lysed as they are with the ΦX174 lysis protein and still absorb at 700nm despite being nonviable. Despite this drawback, ccdB has a natural, specific antitoxin, ccdA, whose sequestering action enables this circuit’s function. Either alternative sequestration methods, like RNA-RNA sequestration, or alternative toxin-antitoxin pairs would be required to recapitulate the design of this circuit and substitute a different toxin.

We collected population measurements using three techniques. Absorbance and fluorescence measurement in an incubator/plate reader gives us the best time resolution for population dynamics, but requires a large number of experimental controls to determine composition and does not accurately measure cell density when using ccdB toxin. Flow cytometry can count the number of cells of each type present in a population, but also suffers from noise in fluorescence detection, especially at the low detection thresholds necessary for processing bacterial samples. Due to this noise, estimates of culture density are not reliably accurate.

CFU counting methods using fluorescent imaging accurately measure both total cell density and composition. High time resolution is labor-intensive to achieve, but the measurement is much more accurate and will be our go-to technique for the future.

The system we designed and tested has two inputs and two control functions; it is natural to expect that each input independently controls a control function, or that simple relationships exist between control functions and the total inducer concentration or the ratio of the inducers. While expressions relating inducer concentrations to total cell density can be obtained analytically, a similar expression for community composition was frustratingly elusive. Addition of AHL degradation to the system will likely simplify our understanding of the relationship between inducers and control functions. Future experiments will test the effects of AHL degradation on the stability and dynamic performance of this population density and composition control circuit.

## IV. MATERIALS AND METHODS

### A. E. coli cell strains

The genome of *E. coli* strain DH5*α*Z1 was integrated with a construct expressing RpaR and CinR using the pOSIP one-step clonetegration kit. The pOSIP plasmid kit used for clonetegration was a gift from Drew Endy and Keith Shearwin (Addgene kit #1000000035) [21]. This strain, DH5*α* Z1-CRp, was used for the first circuit assembly screening presented in (Fig. 2).

The Marionette cell strain used is the “Marionette Wild” strain from Meyer *et. al.* [20]. This cell strain was used in the second circuit assembly screen that yielded A and B cell variants A6, B7 and B8 that were used in the experiments presented in Figures 3 and 4.

DB3.1 ccdB-resistant *E. coli* were used to amplify and purify ccdB containing A1 and B1 plasmids. These cells contain the mutant gyrA462 DNA gyrase, rendering them resistant to ccdB toxicity. DB3.1 cells were obtained from the Belgian Co-ordinated Collections of Microorganisms, accession number LMBP 4098. DB3.1 was originally sold by Invitrogen, but has been discontinued as a product.

Base strains (DH5*α*Z1-CRp or Marionette Wild) were first transformed with A2 or B2 plasmids (see below) that do not contain ccdB toxin, these plasmid containing cells were then transformed a second time with ccdB containing A1 or B1 plasmids purified from DB3.1 cells.

### B. Plasmids and plasmid construction

Each cell line contains 2 plasmids A/B 1 and A/B 2, described below. All plasmids were assembled using the method detailed in *Halleran et al.* [19]. All promoter sequences are taken from *Meyer et al.* [20] to make use of the optimized expression characteristics between the Marionette receptor molecules and their associated evolved promoters. All parts are sourced from the Murray Lab parts library, which is currently in submission to Addgene for distribution. A2 and B2 plasmids contain a ccdA expression unit and an AHL synthase. They replicate using a low-copy ColE1 origin and express kanamycin resistance. The specific constructs are detailed below in the format (promoter - ribosome binding site - CDS - terminator / …)

**pA2 - DH5***α***Z1-CRp**:

pRpa-BCD8-ccdA-L3S3P11(modified) / pLac-BCD12-CinI-ECK120029600

**pB2 - DH5***α***Z1-CRp**:

pCin-BCD8-ccdA-L3S3P11(modified) / pTet-BCD12-RpaI-ECK120029600

**pA2 - Marionette Wild**:

pLuxB-BCD8-ccdA-L3S3P11(modified) / pTac-B0034-CinI-ECK120029600

**pB2 - Marionette Wild**:

pCin-BCD8-ccdA-L3S3P11(modified) / pSalTTC-B0034-LuxI-ECK120029600

A1 and B1 plasmids contain a ccdB expression unit and a constutitively expressed fluorescent tag. They replicate using a low-copy pSC101 origin and express chloramphenicol resistance. These plasmids were assembled using a pool of ribosome binding sites (*ARL*), the Anderson RBS pool, such that cells transformed with the plasmid assembly each contain a different RBS. These unique plasmid variants were initially transformed into DB3.1 *E. coli* to allow amplification of the ccdB containing plasmids without risk of mutation, purified and sequenced, then transformed into Marionette Wild cells containing the appropriate A2 or B2 plasmid.

**pA1 - DH5***α* **Z1-CRp**:

pCin-*ARL*-ccdB-B0015 / J23106-BCD6-BFP-ECK120029600

**pB1 - DH5***α***Z1-CRp**:

pRpa-*ARL*-ccdB-B0015 / J23106-BCD6-YFP-ECK120029600

**pA1 - Marionette Wild**:

pCin-*ARL*-ccdB-B0015 / J23100-BCD6-CFP-L3S3P11(modified)

**pB1 - Marionette Wild**:

pLuxB-*ARL*-ccdB-B0015 / J23100-BCD6-sfYFP-L3S3P11(modified)

### C. Cell growth experiments

*Screening for functioning A and B cell variants* Candidate A cell and a B cell variants were separately grown from picked colonies in LB media overnight, diluted to OD 1, then mixed in all possible combinations in a 1:5 A:B ratio into fresh LB media with half-strength kanamycin (25*µ*g/mL) and chloramphenicol (12.5*µ*g/mL) and aliquoted in triplicate in 500*µ*L into a square 96 well Matriplate (dot Scientific, MGB096-1-1-LG-L). The plate was incubated for 23 hours in a Biotek Synergy H2 incubator/plate reader at 37*?*C while OD600 and fluorescence measurements were taken every 10 minutes. Inducers were added to the 96 well Matriplate before cell suspensions were aliquoted. Induced mixtures were induced with 1mM IPTG and 351*µ*M aTC, while uninduced mixtures received no inducers. A Labcyte Echo 525 Liquid Handler was used to aliquot inducers into each well of the plate before cell suspensions were added.

At hours 0, 7, 19, 23, 10*µ*L of mixed culture in each well was sampled into 15% glycerol and frozen at −80*?*C for community quantification by flow cytometry

*A=B community induction with dilution* An A cell and a B cell variant were separately grown from a frozen glycerol stock in LB media overnight, diluted to OD 1, then mixed in all possible combinations in a 1:1 A:B ratio into fresh LB media with half-strength kanamycin (25*µ*g/mL) and chloramphenicol (12.5*µ*g/mL) and aliquoted in triplicate in 500*µ*L into a square 96 well Matriplate containing inducers pipetted into the plate using the Labcyte Echo. The plate was incubated for 18 hours in a Biotek Synergy H2 incubator/plate reader at 37*?*C while OD600 and fluorescence measurements were taken every 10 minutes. At 18 hours, the plate was removed from the incubator, 90% of the contents of each well was removed, the Labcyte Echo was used to pipet new inducer at each well’s original inducer concentration, and fresh LB medium was added up to 500*µ*L, yielding a 10x culture dilution into identical inducer conditions.

At hours 0, 18, 18 post-dilution, 25 and 43.5, the mixed culture in each well was sampled into 15% glycerol and frozen at −80*?*C for colony counting.

### D. Density and Composition quantification

*Flow cytometry* Frozen cell samples were diluted 30x into PBS buffer containing Syto 62 nuclear stain (Thermo S11344) and incubated on ice for 30 minutes. These samples were then analyzed on a Miltenyi MACSQuant flow cytometer using the mKate/APC channel to detect Syto labeled cells from detector noise, GFP channel to detect YFP and the CFP channel to detect BFP or CFP. FCS files were unpacked to pandas dataframes using the *fcsparser* [22] python package. *Colony counting* Frozen cell samples were diluted 10^4^x into fresh LB media, then 10*µ*L of this suspension was spread on LB agar petri dishes. These plates were incubated at 37*?*C overnight, then colonies were counted. The number of colonies grown was multiplied by the dilution factor to obtain cells/mL.

### E. Modeling and simulations

#### 1) Mathematical model

The model dynamics can be described by the following ordinary differential equations. The description of the model species and the model parameters are given in Tables (I, II) respectively. Note that the subscripts 1 and 2 in the model correspond to the cell strains A and B respectively. Parameter guesses for the inducers, the signals, and the promoter strengths were taken from [20].

**TABLE I.**
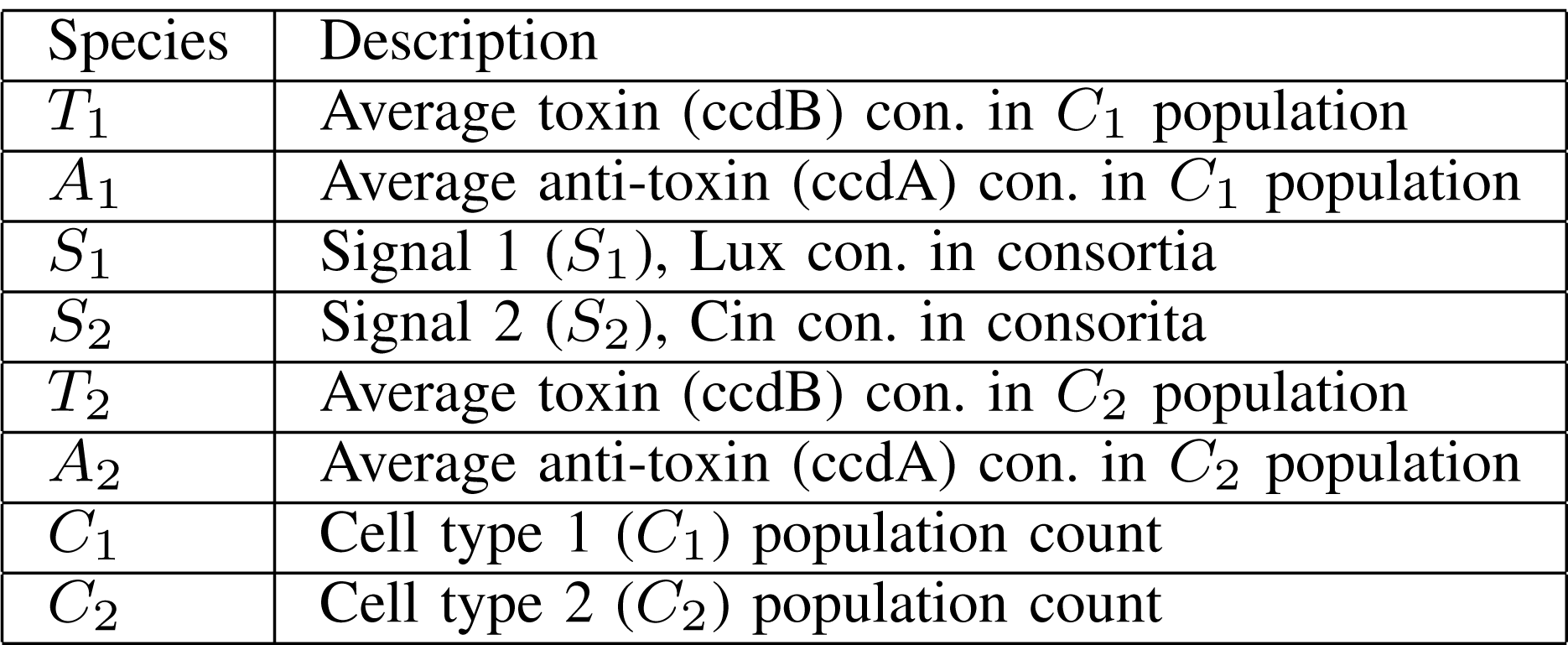
Model species

**TABLE II.** Model parameters

| S.no. | params | description | unit | Guess |
| --- | --- | --- | --- | --- |
| 1 | $\beta_{S_1}$ | Max transcription rate of Lux inducible promoter | con./hr | 6 |
| 2 | $l_{S_1}$ | Leak constant of Lux inducible promoter | N/A | 2e-3 |
| 3 | $K_{S_1}$ | Activation constant of Lux inducible promoter | con. | 430 |
| 4 | $k_b$ | Binding rate between toxin and anti-toxin | 1/con.hr | 30 |
| 5 | $\beta_{S_2}$ | Max transcription rate of Cin inducible promoter | con./hr | 6 |
| 6 | $l_{S_2}$ | Leak constant of Cin inducible promoter | N/A | 2e-3 |
| 7 | $K_{S_2}$ | Activation constant of Cin inducible promoter | con. | 30 |
| 8 | $\beta_{tac}$ | Max transcription rate of inducible promoter $P_{tac}$ | con./hr | 19.8e-3 |
| 9 | $l_{tac}$ | Leak constant of inducible promoter $P_{tac}$ | N/A | 1.5e-3 |
| 10 | $K_{tac}$ | Activation constant of inducible promoter $P_{tac}$ | con. | 1.4e5 |
| 11 | $\beta_{sal}$ | Max transcription rate of inducible promoter $P_{sal}$ | con./hr | 14.4e-3 |
| 12 | $l_{sal}$ | Leak constant of inducible promoter $P_{sal}$ | N/A | 2.1e-4 |
| 13 | $K_{sal}$ | Activation constant of inducible promoter $P_{sal}$ | con. | 4.3e4 |
| 14 | $k_C$ | Cell Division rate | 1/hr | 0.6 |
| 15 | $C_{max}$ | Population cap | conc. | 5500 |
| 16 | $d_c$ | Cell death rate | 1/conc.hr | 0.8 |
| 17 | $S_2$ | Max induced Cin concentration | con. | 100 |
| 18 | $I$ | Max induced IPTG concentration | con. | 1e6 |
| 19 | $S_1$ | Max induced Lux concentration | con. | 1e3 |
| 20 | $Sal$ | Max induced Sal concentration | con. | 1e5 |
| 21 | $K_{tox}$ | Repression coefficient of toxin to proliferation | con. | 5 |
| 22 | $d_S$ | Degradation constant of AHLs | 1/hr | 0.1 |
| 23 | $d$ | Basal degradation of each cell | 1/hr | 0.01 |
| 24 | $d_T$ | Basal degradation of toxins and antitoxins | 1/hr | 0.01 |
| 25 | $K_r$ | Ribosome scaling factor | N/A | 5 |

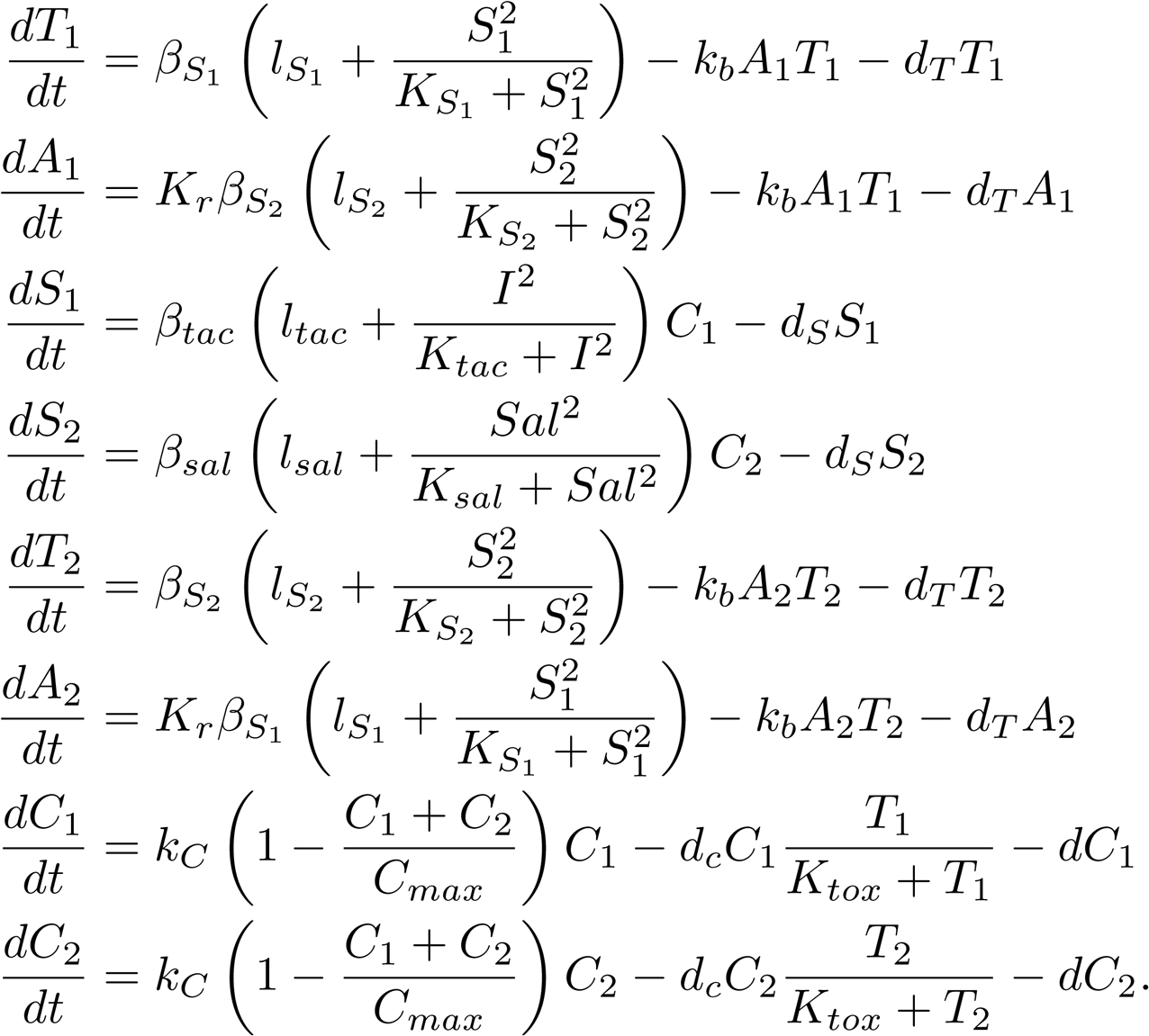

#### 2) Simulations

All simulations of the ODE model were performed using the Python SciPy library [23]. The code to regenerate the simulations is available at [24].

#### 3) Sensitivity analysis

We performed sensitivity analysis of the model to identify the parameters that are most sensitive in the model. The sensitivity analysis results are shown in Fig. 6. It is clear that one of the most sensitive parameters is *β*_*S*2_ — the maximum transcription rate of Cin inducible promoter that drives the production of the toxin ccdB. We used these inferences from the sensitivity analysis to guide the screening experiments of the ribosome binding site driving the ccdB production as discussed in Section II.

**Fig. 6.**
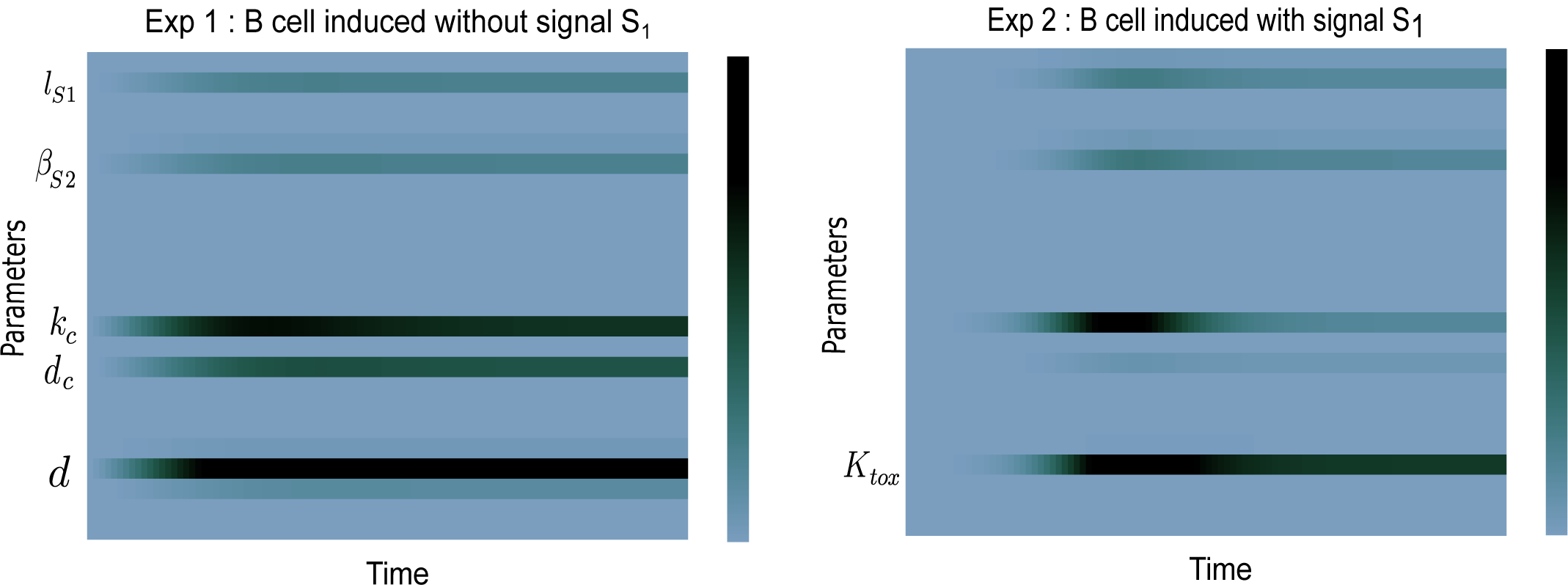
Sensitivity analysis of the mathematical model of this system reveals significant sensitivity to a few model parameters, particularly 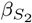, the production rate of ccdB; *k*_*c*_, the cell division rate; *d*_*c*_, the cell death rate. The only tunable parameter of these is 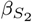, which is modified in our parameter screen for functional community circuits.

## ACKNOWLEDGMENT

The authors would like to thank Chelsea Hu, Leopold Green, Andrey Shur and Mark Prator for discussion and assistance in performing computational and laboratory work. The authors R. D. M. and A. P. are supported by Defense Advanced Research Projects Agency (Agreement HR0011-17-2-0008). The content of the information does not necessarily reflect the position or the policy of the Government, and no official endorsement should be inferred.

